# RUNX3 regulates cell cycle-dependent chromatin dynamics by functioning as a pioneer factor of the restriction point

**DOI:** 10.1101/409227

**Authors:** Jung-Won Lee, Da-Mi Kim, Ju-Won Jang, Tae-Geun Park, You-Soub Lee, Xin-Zi Chi, Il Yeong Park, Jin-Won Hyun, Yoshiaki Ito, Suk-Chul Bae

**Affiliations:** Department of Biochemistry, School of Medicine, and Institute for Tumor Research, Chungbuk National University, Cheongju, 28644, South Korea; College of Pharmacy, Chungbuk National University, Cheongju, 361-763, South Korea; Department of Biochemistry, School of Medicine, Jeju National University, Jeju, 63243, South Korea; Cancer Science Institute of Singapore, National University of Singapore, 14 Medical Drive, Singapore 117599

**Keywords:** RUNX3, K-Ras, ARF, p53, restriction point, chromatin dynamics, pioneer factor

## Abstract

The cellular decision regarding whether to undergo proliferation or death is made at the restriction (R)-point, which is disrupted in nearly all tumors. The identity of the molecular mechanisms that govern the R-point decision is one of the fundamental issues in cell biology. We found that early after mitogenic stimulation, RUNX3 bound to its target loci, where it opened chromatin structure by sequential recruitment of Trithorax group proteins and cell-cycle regulators to drive cells to the R-point. Soon after, RUNX3 closed these loci by recruiting Polycomb repressor complexes, causing the cell to pass through the R-point toward S phase. If the RAS signal was constitutively activated, RUNX3 inhibited cell cycle progression by maintaining R-point-associated genes in an open structure. Our results identify RUNX3 as a pioneer factor for the R-point and reveal the molecular mechanisms by which appropriate chromatin modifiers are selectively recruited to target loci for appropriate R-point decisions.

## Introduction

In response to mitogenic stimulation, the cell makes a critical decision regarding whether to advance into late G_1_, retreat into G_0_, or undergo apoptosis. This decision occurs at the restriction (R)-point, and the associated decision-making machinery is perturbed in nearly all cancer cells ^1–3^. The transition is governed by R-point–associated proteins, including pRB, E2F, Cyclins, CDKs, p21, p27, and c-Myc, with pRB serving as the primary molecular regulator ^3^. The R-point decision is made after Cyclin D-CDK4/6 activation and before Cyclin E-CDK2 activation ^3^. Although the R-point decision is literally a matter of life and death for the cell and is critical to the health of the organism, it remains unclear how each cell chooses its own fate.

The R-point decision-making process involves regulation of several hundred genes ^4^. For silent genes to be induced, target sites within their regulatory regions must be bound *de novo* by transcription factors, which initiate their expression. The special transcription factors that have the capacity to associate with condensed chromatin independently of other factors and modulate chromatin accessibility are known as pioneer factors ^5^. Proteins of several families possess these properties, including FOXA (also known as HNF3) ^6^, OCT4, SOX2, and KLF ^7^. Pioneer factors usually open chromatin, but can close chromatin in some cases, e.g., FOXA1 recruits TLE to make chromatin inaccessible ^8^.

To modulate chromatin accessibility and regulate gene transcription, pioneer factors require a complex network of other proteins, including coactivators, corepressors, histone-modifying complexes, chromatin-remodeling complexes, and the basal transcription machinery. For example, the Trithorax group (TrxG) and Polycomb group (PcG) proteins establish histone modifications that activate and repress transcription, respectively. TrxG proteins can be broadly classified into two categories: histone modifiers ^9^ and nucleosome remodelers ^10^. The TrxG histone modifiers include mixed-lineage leukemia family members (MLLs), which methylate H3 at lysine 4 (H3K4-me3, -me2, and -me1), a mark that favors transcriptional activation. On the other hand, the TrxG nucleosome remodelers include the SWI–SNF complex, which facilitates binding of transcription factors and the basal transcription machinery. PcG complexes are classified into two categories: Polycomb repressor complex 1 and 2 (PRC1 and PRC2). Both complexes consist of multiple proteins: PRC1 contains BMI1 and ring finger protein 1 (RING1) or ring finger protein 2 (RNF2) ^11^, whereas PRC2 contains EED and enhancer of zeste homologs (EZH1 and EZH2) that trimethylate H3 at lysine 27 (H3K27-me3), a characteristic of inactive chromatin ^12^. Via recruitment of these chromatin modulators, cells regulate signal-dependent gene expression at the correct target loci at the right time. The underlying mechanism, which represents one of the most fundamental issues in molecular biology, remains poorly understood.

The DNA-binding transcription factor RUNX3, which plays pivotal roles in lineage determination, defends against oncogenic K-RAS–induced tumorigenesis ^13^. Deletion of *Runx3* in mouse lung results in development of lung adenomas and accelerates K-Ras–induced progression into adenocarcinomas (ADCs) ^14^. In mouse embryonic fibroblasts, *Runx3* deletion perturbs the R-point, leading to transformation ^4^. Here, we demonstrate that RUNX3 is a pioneer factor of the R-point that plays a key role in sequential recruitment of TrxG and PcG proteins to target loci in a RAS signal-dependent manner, enabling an appropriate R-point decision.

## Results

### The RUNX3–BRD2–nucleosome complex recruits SWI/SNF and TFIID before the R-point

The R-point decision is made 3–4 hr after serum stimulation ^15^. Previously, we showed that the RUNX3–BRD2 complex forms 1–2 hr after serum stimulation ^14^, and that this complex contributes to the R-point decision by regulating hundreds of genes ^4^. An important question regarding the R-point is how the chromatin structure of target genes is selectively activated in a signal-dependent manner. We obtained a clue about this issue from our previous observation that p300 histone acetyltransferase acetylates not only histones but also RUNX3 ^14^. In addition, BRD2 contains two bromodomains (BD1 and BD2), each of which interacts with a distinct protein: BD1 binds RUNX3 acetylated at Lys-94 and Lys-171 ^14^, whereas BD2 binds the acetylated histones H4K5-ac, H4K12-ac, and H3K14-ac ^16,17^ (Fig. 1a). Notably, we detected interactions between p300, RUNX3, and H4K12-ac 1–2 hr after mitogenic stimulation (Fig. 1b, RX3/p300, RX3/H4K12-ac), as well as between BRD2, RUNX3, and H4K12-ac (Fig. 1b, BRD2/RX3 and BRD2/H4K12-ac). The RUNX3–H4K12-ac interaction was markedly diminished by knockdown of *BRD2* (see below). These results suggest that RUNX3 guides p300 to target loci, where it acetylates histones, and that BRD2 binds both acetylated RUNX3 and acetylated histones through its two bromodomains, prior to the R-point.

**Fig. 1:**
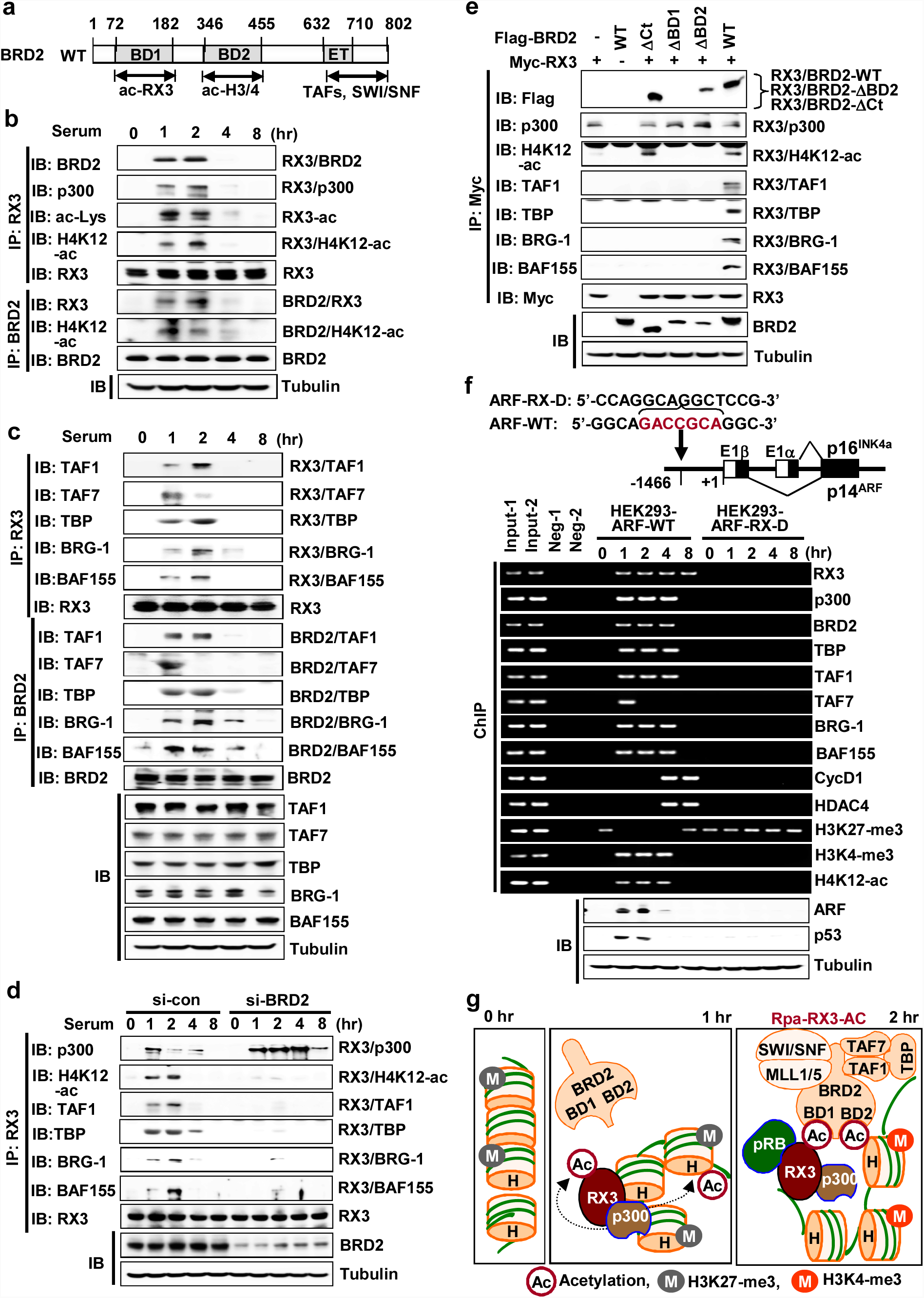
The RUNX3–BRD2–nucleosome complex recruits SWI/SNF and TFIID before the R-point. **a** Schematic diagram of BRD2 structure and interacting proteins. BD1 interacts with RUNX3 acetylated at Lys-94 and Lys-171; BD2 interacts with acetylated histones H4K4-ac, H4K12-ac, and H3K14-ac; and the C-terminal region interacts with the TFIID and SWI/SNF complexes. **b-c** HEK293 cells were serum-starved for 24 hr, and then stimulated with 10% serum. Cells were harvested at the indicated time points, and the levels of acetylated RUNX3 (RX3-ac), TAF1, TAF7, TBP, BRG-1, and BAF155 were measured by IP and IB. The time-dependent interactions between RUNX3, BRD2, p300, H4K12-ac, TFIID complex (TAF1, TAF7, and TBP), and SWI/SNF complex (BRG-1 and BAF155) were measured by IP and IB. **d** HEK293 cells were treated with control siRNA (si-con) or BRD2-specific siRNA (si-BRD2), serum-starved for 24 hr, and then stimulated with 10% serum for the indicated durations. The time-dependent interactions between RUNX3, p300, H4K12-ac, TFIID complex (TAF1, TAF7, and TBP), and SWI/SNF complex (BRG-1 and BAF155) were measured by IP and IB. **e** HEK293 cells were transfected with Myc-RUNX3, Flag-BRD2-WT, Flag-BRD2-ΔCt (lacking C-terminal aa 633–802), Flag-BRD2-ΔBD1 (lacking BD1), or Flag-BRD2-ΔBD2 (lacking BD2). Cells were serum-starved for 24 hr, and then stimulated with 10% serum. Cells were harvested after 2 hr, and the interactions of Myc-RUNX3 with p300, BRD2, H4K12-ac, TFIID complex (TAF1 and TBP), and SWI/SNF complex (BRG-1 and BAF155) were measured by IP and IB. **f** The RUNX3-binding site (GACCGCA) in the *ARF* enhancer region (ntd −1466) was deleted in HEK293 cells by the CRISPR/Cas9 method to obtain the HEK293-ARF-RX-D cell line. Deletion of the RUNX3-binding site was confirmed by nucleotide sequencing. Wild-type HEK293 cells (HEK293-ARF-WT) and HEK293-ARF-RX-D cells were serum-starved for 24 hr. The cells were then treated with 10% serum, and the binding of RUNX3, p300, BRD2, TFIID complex (TAF1, TAF7, and TBP), SWI/SNF complex (BRG-1 and BAF155), Cyclin D1, HDAC4, H3K27-me3, H3K4-me3, and H4K12-ac to the *ARF* promoter was measured by ChIP at the indicated time points. One-thirtieth of the lysates were PCR-amplified as input samples. Time-dependent expression of *ARF* and *p53* was measured by IB. **g** Schematic illustration of sequential molecular events at RUNX3 target loci during R-point regulation. RUNX3 binds to condensed chromatin marked by H3K27-me3 (inhibitory mark). p300 recruited to the loci acetylates RUNX3 and histones. Then, BRD2 binds both acetylated RUNX3 and acetylated histone through its two bromodomains. At 1 hr after serum stimulation, SWI/SNF and TFIID are recruited to the loci through the C-terminal region of BRD2 to form Rpa-RX3-AC, and H3K27-me3 is replaced by H3K4-me3 (activating mark).

BRD2 interacts with the SWI/SNF and TFIID complexes through its C-terminal region ^17,18^ (Fig. 1a), suggesting that RUNX3 interacts with these complexes through BRD2. Analysis of the interactions of RUNX3 and BRD2 with the components of these complexes at various time points after serum stimulation revealed that TAF1 (activating TAF), TAF7 (inhibitory TAF), and TBP formed a complex with BRD2 and RUNX3 1 hr after mitogenic stimulation (Fig. 1c). Soon thereafter, TAF7 dissociated from the complex (Fig. 1c), suggesting that TFIID is activated after the interaction with RUNX3– BRD2. After 4 hr, TAF1 and TBP also dissociated from RUNX3 (Fig. 1c). Similarly, BRG-1 and BAF155 (components of the SWI/SNF complex) also interacted with RUNX3 and BRD2 1–2 hr after mitogenic stimulation and dissociated at 4 hr (Fig. 1c). The interactions of SWI/SNF and TFIID complexes with the RUNX3–BRD2 complex were confirmed by the proximity ligation assay (PLA) (Supplementary Fig. 1a). Consistently, expression of R-point–associated proteins [p14^ARF^ (hereafter ARF), p53, and p21] was induced at the same time that RUNX3 interacted with BRD2, SWI/SNF, and TFIID (Supplementary Fig. 1b).

The interactions of RUNX3 with H4K12-ac and the components of TFIID and SWI/SNF were markedly weakened by knockdown of *BRD2*, indicating that H4K12-ac and the TFIID and SWI/SNF complexes interact with RUNX3 through BRD2 (Fig. 1d). The requirement of BRD2 for these interactions was further confirmed by transfection of WT and RUNX3 mutants with BRD2. SWI/SNF and TFIID interacted with RUNX3 lacking the N-terminal 53 amino acid (aa) region (RX3-ΔNt), but not with a RUNX3 mutant lacking Lys residues critical for the interaction with BRD2 (RX3-KR-94-171) (Supplementary Fig. 1c). RUNX1, which shares high sequence similarity with RUNX3, directly interacts with TAF1 though an interaction between acetylated Lys-43 of RUNX1 and the bromodomain of TAF1 ^19^. However, RUNX3 does not contain a lysine residue corresponding to Lys-43 of RUNX1 (Supplementary Fig. 1d), and therefore requires BRD2 for its interaction with SWI/SNF and TFIID.

### RUNX3 is a pioneer factor of the R-point

To elucidate the temporal order of the molecular interactions among RUNX3, BRD2, histones, SWI/SNF, and TFIID, we transfected cells with Myc-RUNX3 along with Flag-BRD2-WT, Flag-BRD2-ΔCt (lacking aa 633–802 in the C-terminal region), Flag-BRD2-ΔBD1 (lacking BD1), or Flag-BRD2-ΔBD2 (lacking BD2), and then performed immunoprecipitation/immunoblotting (IP/IB) to analyze their interactions 2 hr after serum stimulation. Because the RUNX3–p300 interaction is independent of BRD2 (Fig. 1d), it was not affected by expression of mutant BRD2 (Fig. 1e). Therefore, the RUNX3–p300 interaction must occur earlier than the RUNX–BRD2 interaction. Deletion of the C-terminal region of BRD2 (Flag-BRD2-ΔCt) abolished the interactions of RUNX3 with TFIID and SWI/SNF without affecting the RUNX3–H4K12-ac interaction (Fig. 1e), whereas deletion of either BD1 or BD2 abolished the RUNX3– H4K12-ac interaction (Fig. 1e). These results confirmed that BRD2 simultaneously binds acetylated RUNX3 and acetylated H4 through BD1 and BD2, respectively. Notably, however, BRD2-ΔBD2 failed to bridge between RUNX3, SWI/SNF, and TFIID (Fig. 1e). These results suggest that only BRD2 bound to both RUNX3 and histone (RUNX3–BRD2–nucleosome complex) can interact with the SWI/SNF and TFIID complexes. On the basis of these findings, we conclude that RUNX3 on its binding site in an enhancer region facilitates p300-mediated histone acetylation around the locus, juxtaposes the promoter region via formation of the RUNX3–BRD2–H4K12-ac complex, and then recruits SWI/SNF and TFIID complexes to the locus via BRD2.

To confirm RUNX3-mediated recruitment of SWI/SNF and TFIID complexes to specific chromatin loci, we chose the *ARF* (*CDKN2A*) locus as a model. *ARF* is a target of RUNX3 ^14^ that is critical for the life and death of cells; thus, regulation of its expression could represent the R-point decision. The *ARF* promoter contains a perfect match to the RUNX consensus binding site 1466 bases upstream of the transcription initiation site ^20^. Using the CRISPR/Cas9 system, we deleted the RUNX-binding site in HEK293 cells to obtain HEK293-ARF-RX-D cells. Chromatin IP (ChIP) analysis revealed that in parental HEK293 cells (HEK293-ARF-WT), RUNX3 bound to the *ARF* locus 1 hr after serum stimulation, and this interaction was maintained for 8 hr (Fig. 1e). p300, BRD2, and components of SWI/SNF and TFIID were recruited to the locus 1 hr after serum treatment, and the interaction was maintained for 4 hr (except for TAF7, which dissociated after 2 hr) (Fig. 1e). By contrast, in HEK293-ARF-RX-D cells, none of these proteins was recruited to the locus (Fig. 1f), indicating that binding of RUNX3 to the RUNX consensus site is critical for recruitment of the chromatin-remodeling complex and basal transcriptional machinery to its target locus.

In HEK293-ARF-WT cells, H3K27-me3 (a repressive histone modification) was enriched within the *ARF* promoter region prior to serum stimulation (0 hr) (Fig. 1f). Notably, H3K27-me3 was replaced by H3K4-me3 (an activating histone modification) 1–4 hr after serum stimulation, whereas H3K27-me3 was restored 8 hr later (Fig. 1f). Similarly, H4K12-ac (an activating histone modification) was detected 1–4 hr after stimulation, but then disappeared (Fig. 1f). By contrast, in HEK293-ARF-RX-D cells, the H3K27-me3 modification was maintained, and H4K12 was not acetylated for a long time after serum stimulation (Fig. 1f). Consistently, induction of ARF leading to p53 stabilization was detected 1–2 hr after stimulation in HEK293-ARF-WT cells, but not in HEK293-ARF-RX-D cells (Fig. 1f). These results demonstrate that RUNX3 associates with its binding site and opens and activates the chromatin structure of target loci 1–2 hr after serum stimulation, and then subsequently closes the loci, indicating that RUNX3 is a pioneer factor of the R-point. We named the RUNX3-containing complex formed before the R-point as the R-point–associated RUNX3-containing activator complex (Rpa-RX3-AC) (Fig. 1g).

In HEK293-ARF-WT cells, BRD2, SWI/SNF, and TFIID continued to be associated with the *ARF* locus up to 4 hr after stimulation (Fig. 1f), but these proteins interacted with RUNX3 for only up to 2 hr (Fig. 1b, c). These results suggest that BRD2 remains bound to the *ARF* locus for a while even after dissociation from RUNX3, most likely through interactions with histones and other DNA-binding proteins recruited to open chromatin. The sequence of molecular events involved in binding of RUNX3 to chromatin, recruitment of SWI/SNF and TFIID, and alterations of histone marks are summarized in Fig. 1g. This model suggests that the enhancer interacts with the promoter through Rpa-RX3-AC during the R-point (Fig. 1g). The enhancer–promoter interaction on the *ARF* locus 2 hr after serum stimulation was confirmed by the chromosome conformation capture assay (3C assay ^21^) (Supplementary Fig. 2).

**Fig. 2:**
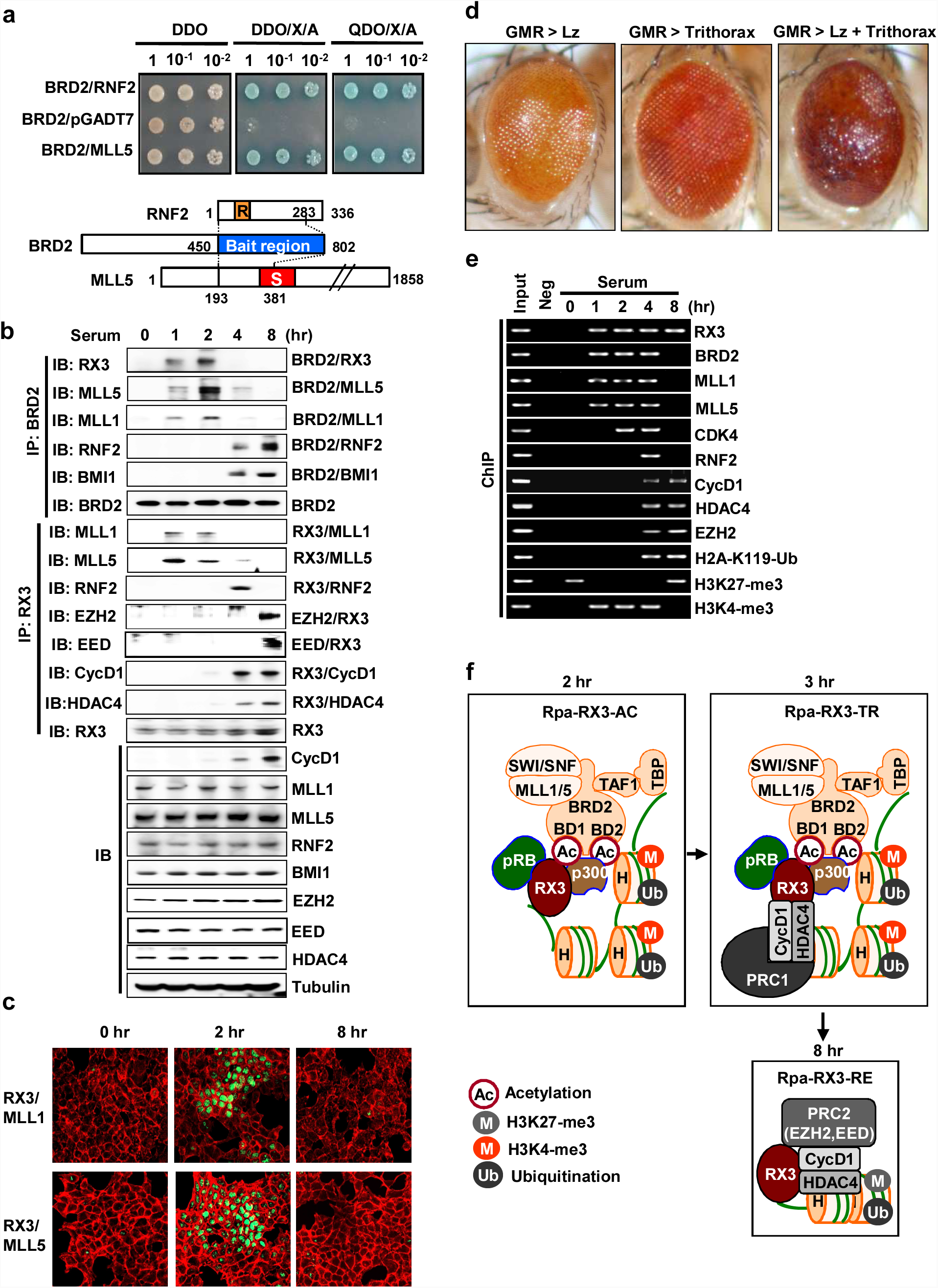
RUNX3 sequentially recruits TrxG and PcG complexes before and at the R-point to form a large complex. **a** Yeast two-hybrid screening using Gal4-BRD2 (aa 450–802) as bait identified RNF2 and MLL5 as BRD2-binding proteins (see STAR methods). DDO=SD-Leu/-Trp, DDO/X/A=SD-Leu/-Trp/X-a-gal/ABA, QDO/X/A=SD-Leu/-Trp/-His/-Ade/X-a-gal/ ABA medium. Selective colonies were identified by DNA sequencing. **b** HEK293 cells were serum-starved for 24 hr, and then stimulated with 10% serum. Cells were harvested at the indicated time points, and the time-dependent interactions between RUNX3, BRD2, Cyclin D1, MLL1, MLL5, RNF2, BMI1, EZH2, EED, and HDAC4 were measured by IP and IB. **c** PLA showing RUNX3-MLL1 and RUNX3-MLL5 at the indicated time points after serum stimulation. Green fluorescence indicates association of the indicated proteins. F-actin was stained (red) to visualize the cytoplasmic compartment. **d** Microscopy images of transgenic fly eyes. *Lozenge* is a *Drosophila* homolog of the *RUNX* genes. *Glass multimer reporter* (*GMR*)*-Gal4* promotes eye-specific expression of *UAS*-inserted genes. *GMR*-driven *Lozenge* overexpression (*GMR*-*Gal4/+;UAS-Lozenge* → GMR>Lozenge) or *GMR*-driven *Trithorax* overexpression (*GMR*-*Gal4/+;Trx*G14137 → GMR>Trithorax) conferred weak rough phenotypes. However, *GMR-*driven overexpression of both *Lozenge* and *Trithorax* (*GMR>Lozenge+Trithorax*) resulted in a severe defective eye phenotype with loss of external ommatidial facets. **e** HEK293 cells were serum-starved for 24 hr, and then stimulated with 10% serum. The binding of RUNX3, BRD2, MLL1, MLL5, CDK4, RNF2, Cyclin D1, HDAC4, EZH2, H2A-K119-Ub, H3K27-me3, and H3K4-me3 to the *ARF* promoter was measured by ChIP at the indicated time points. One-thirtieth of the lysates were PCR-amplified as input samples. **f** Schematic illustration of the R-point transition. At 1 hr after serum stimulation, RUNX3 associates with various proteins, including p300, BRD2, H4K12-ac, SWI/SNF, TFIID, and MLL1/5, to form Rpa-RX3-AC. Between 2 and 4 hr after serum stimulation, Rpa-RX3-AC interacts with PRC1–CyclinD1–HDAC4 to from a transient complex, Rpa-RX3-TR. Subsequently, Rpa-RX3-TR is destroyed (at 4 hr) to form Rpa-RX3-RE (at 8 hr).

### RUNX3 also recruits MLL1/5, PRC1, and PRC2 at different stages of the R-point

To understand how the RUNX3–BRD2 complex controls histone modifications, we performed yeast two-hybrid screening using the C-terminal region of BRD2 (aa 450–802) as bait. The screen identified MLL5 and RNF2, which play opposing roles in chromatin dynamics (Fig. 2a): MLL5 is a TrxG histone modifier that contributes to chromatin activation, whereas RNF2 is a component of the PRC1 complex, which inactivates chromatin.

Analysis of the interactions among BRD2 and MLL family members revealed that BRD2 interacted with MLL1 and MLL5 1–2 hr after serum stimulation (Fig. 2b). These proteins also interacted with RUNX3 at the same time points (Fig. 2b). The physical interactions between MLL1/5 and RUNX3 at 2 hr after serum stimulation were confirmed by the PLA (Fig. 2c). A genetic interaction between *Lozenge* (a *Drosophila* homolog of the *RUNX* family genes) and *Trithorax* (a *Drosophila* homolog of the *MLL* family genes) was also observed during fly eye development (Fig. 2d). ChIP analysis revealed that MLL1/5 bound to the *ARF* locus 1–4 hr after stimulation, and H3K4-me3 was enriched at the locus at these time points (Fig. 2e). These results suggest that MLL1/5 recruited to chromatin through the interaction with the RUNX3–BRD2 complex forms activating histone modifications. Therefore, MLL1/5 are additional components of Rpa-RX3-AC.

Analysis of interactions among BRD2 and components of PRC1 revealed that BRD2 interacted with RNF2 and BMI1 4–8 hr after serum stimulation (Fig. 2b). At that time, RUNX3 associated with Cyclin D1 and HDAC4 instead of BRD2 (Fig. 2b). EED and EZH2 (components of PRC2) associated with RUNX3 8 hr after stimulation (Fig. 2b). ChIP analysis revealed that Cyclin D1, HDAC4, and EZH2 were bound to chromatin after the R-point, at a time when H4K12-ac was absent and H3K27-me3 was enriched at the locus (Fig. 2e). After 4 hr, RNF2 briefly interacted with RUNX3 and BRD2 (Fig. 2b) and was recruited to the target locus (Fig. 2e), suggesting that PRC1 and Rpa-RX3-AC form a transient complex immediately before the latter complex is destroyed (about 3 hr after mitogen stimulation). We named the transient assembly as the R-point–associated RUNX3-containing transient complex (Rpa-RX3-TR) (Fig. 2f).

At 4–8 hr after mitogen stimulation, RUNX3 and BRD2 existed in separate complexes: RUNX3 formed a complex with Cyclin D1, HDAC4, and PRC2 (Fig. 2b), which remained bound to target chromatin loci (Fig. 2e), whereas BRD2 formed the BRD2–PRC1 complex (Fig. 2b), which was released from the loci (Fig. 2e). Because the RUNX3–Cyclin D1–HDAC4–PRC2 complex inactivates chromatin, we named it the R-point–associated RUNX3-containing repressor complex (Rpa-RX3-RE) (Fig. 2f). We refer to the Rpa-RX3-AC → Rpa-RX3-TR → Rpa-RX3-RE transition as the R-point transition.

### The PRC1–Cyclin D1–HDAC4 complex interacts with Rpa-RX3-AC to form Rpa-RX3-TR

We next investigated how Rpa-RX3-TR is formed. IP/IB analysis revealed that the RNF2–Cyclin D1 interaction occurred 2 hr after serum stimulation and gradually weakened thereafter (Fig. 3a). The RNF2–HDAC4 interaction was detected only 4 hr after stimulation (Fig. 3a), whereas the HDAC4–Cyclin D1 interaction occurred 4 hr after simulation, and gradually strengthened thereafter (Fig. 3a). These results suggest that PRC1 interacts with Cyclin D1 2 hr after stimulation and subsequently matures to the PRC1–Cyclin D1–HDAC4 complex. Formation of the PRC1–Cyclin D1–HDAC4 complex 4 hr after stimulation was confirmed by transfection followed by IP/IB analysis (Fig. 3b), and the direct interactions between the proteins were confirmed by IP/IB with *in vitro* translated proteins (Fig. 3c, d). Domain mapping analysis revealed that the N-terminal region (aa 1–44) of Cyclin D1 (which interacts with pRB) was required for the interactions with RUNX3 and HDAC4, whereas the C-terminal region (aa 156–295) was required for the interaction with RNF2 (Fig. 3e–h). RUNX3 interacts with Cyclin D1 through its N-terminal region (aa 1–186 of RUNX3) (Supplementary Fig. 3).

**Fig. 3:**
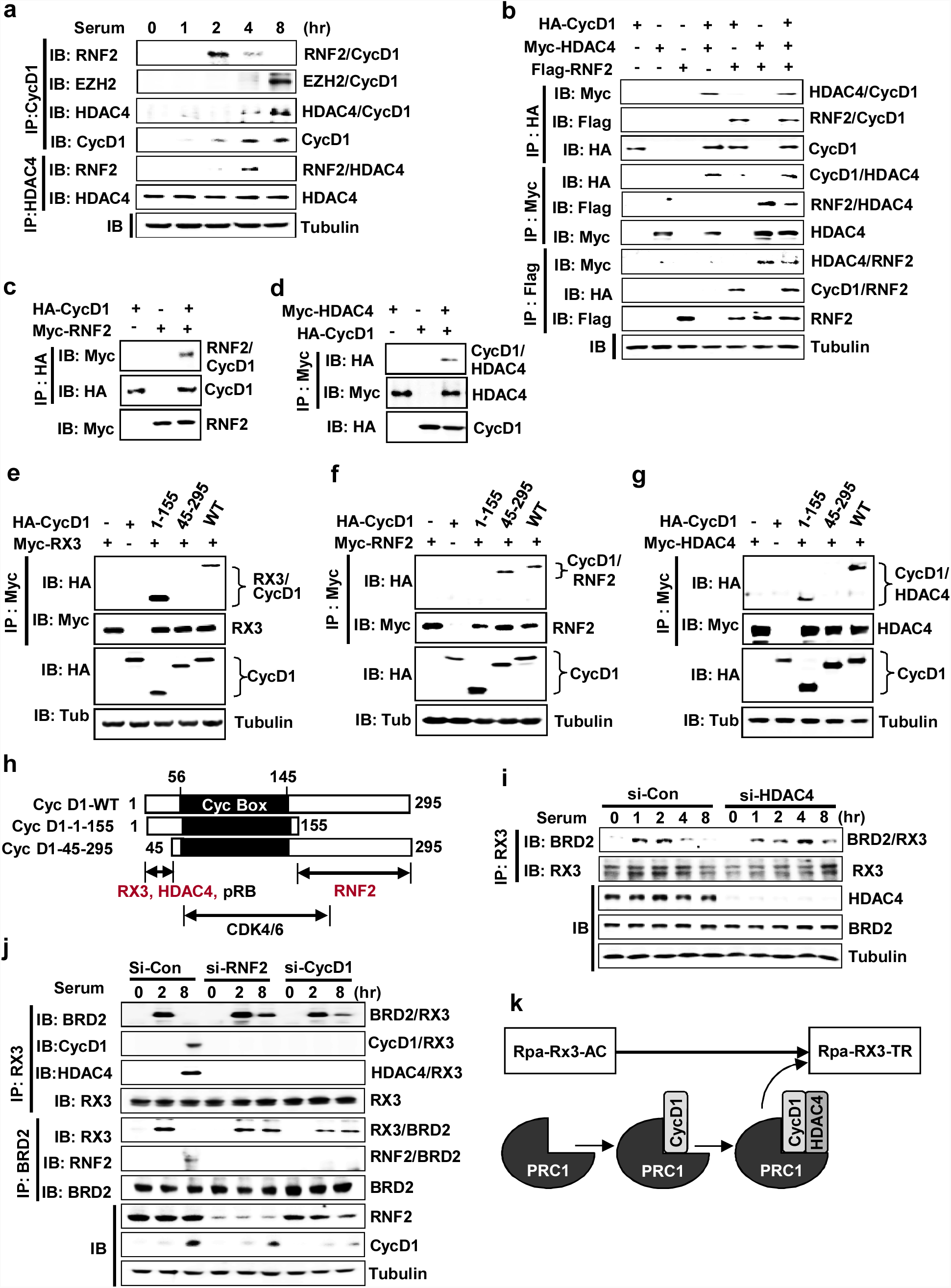
Formation of the PRC1–CyclinD1–HDAC4 complex. **a** HEK293 cells were serum-starved for 24 hr, and then stimulated with 10% serum. Cells were harvested at the indicated time points. Time-dependent formation of the RNF2–Cyclin D1, EZH2–Cyclin D1, HDAC4–Cyclin D1, and RNF2–HDAC4 complexes was measured by IP and IB. **b** HEK293 cells were transfected with HA-Cyclin D1, Myc-HDAC4, and Flag-RNF2, and the interactions between the proteins were measured by IP and IB. **c-d** HA-Cyclin D1, Myc-RNF2, and Myc-HDAC4 were translated *in vitro* and the interactions among the proteins were measured by IP and IB. **e-g** Myc-RUNX3, Myc-RNF2, or Myc-HDAC4 was co-expressed with HA-Cyclin D1(WT), HA-Cyclin D1(1–155), and HA-Cyclin D1(45–295) in HEK293 cells. The interactions of RUNX3, RNF2, and HDAC4 with Cyclin D1 were measured by IP and IB 4 hr after serum stimulation. **h** Regions of Cyclin D1 required for the interaction with RUNX3, RNF2, and HDAC4 are summarized. Cyclin D1 regions known to interact with pRB and CDK4/6 are also indicated. Cyc Box = Cyclin Box. **i** HEK293 cells were treated with control or HDAC4-specific siRNA (si-con or si-HDAC4), serum-starved for 24 hr, and then stimulated with serum for the indicated durations. Time-dependent formation of the BRD2–RUNX3 complex was measured by IP and IB. **j** HEK293 cells were treated with control, RNF2-specific, or Cyclin D1–specific siRNA (si-con, si-RNF2, or si-CycD1), serum-starved for 24 hr, and then stimulated with serum for the indicated durations. Time-dependent formation of the BRD2–RUNX3, Cyclin D1–RUNX3, HDAC4–RUNX3, and RNF2–BRD2 complexes was measured by IP and IB. **k** Schematic illustration of the process of Rpa-RX3-TR formation. Cyclin D1, which is induced 2 hr after serum stimulation, interacts with PRC1 (containing RNF2) and matures into the PRC1–CyclinD1–HDAC4 complex. The PRC1–CyclinD1–HDAC4 complex then interacts with Rpa-RX3-AC to form Rpa-RX3-TR.

PRC1, Cyclin D1, and HDAC4 were simultaneously recruited to target loci 4 hr after serum stimulation (Fig. 2e). Notably, siRNA-mediated knockdown of *HDAC4* inhibited dissociation of RUNX3–BRD2 (Fig. 3i). Similarly, knockdown of either *RNF2* or *CCND1* (the gene that encodes Cyclin D1) diminished the RUNX3–Cyclin D1, RUNX3–HDAC4, and RNF2–BRD2 interactions and effectively inhibited dissociation of RUNX3–BRD2 (Fig. 3j). These results suggest that the PRC1–Cyclin D1–HDAC4 complex, rather than any of the individual components, interacts with Rpa-RX3-AC to form Rpa-RX3-TR (Fig. 3k).

Inactivation of chromatin is associated with HDAC-mediated histone deacetylation and RNF2-mediated H2A ubiquitination at Lys-119 (H2A-K119-Ub) ^11^. Consistently, H4K12 acetylation was reduced (Fig. 1f) and H2A ubiquitination at Lys-119 (H2A-K119-Ub) was enriched at the *ARF* locus 4–8 hr after stimulation (Fig. 2e). These results demonstrate that the PRC1–Cyclin D1–HDAC4 complex binds to Rpa-RX3-AC, forms Rpa-RX3-TR, and then contributes to inactivation of chromatin at target loci by deacetylating H4K12 and ubiquitinating H2A.

### Association of CDK4 and Cyclin D1 leads to formation of Rpa-RX3-TR

Since R-point decision is made after the association of Cyclin D1 and CDK4, we investigated the role of Cyclin D1–CDK4 in the R-point transition. Previously, we showed that hypo-phosphorylated pRB interacts with RUNX3–BRD2 and contributes to R-point commitment ^4^. E2F1 (which associates with hypo-phosphorylated pRB), as well as pRB itself, interacted with RUNX3 1–2 hr after stimulation (Fig. 4a). The pRB–E2F1 complex was released from RUNX3 when pRB was phosphorylated at Ser-795 by Cyclin D1–CDK4/6 (Fig. 4a). These results suggest that CDK4 approaches Rpa-RX3-AC to phosphorylate pRB. IP/IB analysis revealed that CDK4 interacted with RUNX3 2 hr afterward, and that the interaction weakened thereafter (Fig. 4a). CDK4 was also bound to the *ARF* locus while it interacted with RUNX3 (Fig. 2e). The physical interaction between RUNX3 and CDK4 2 hr after serum stimulation was confirmed by the PLA (Fig. 4b). These results demonstrate that CDK4, along with pRB and E2F1, which play key roles in cell-cycle regulation, is a component of Rpa-RX3-AC.

**Fig. 4:**
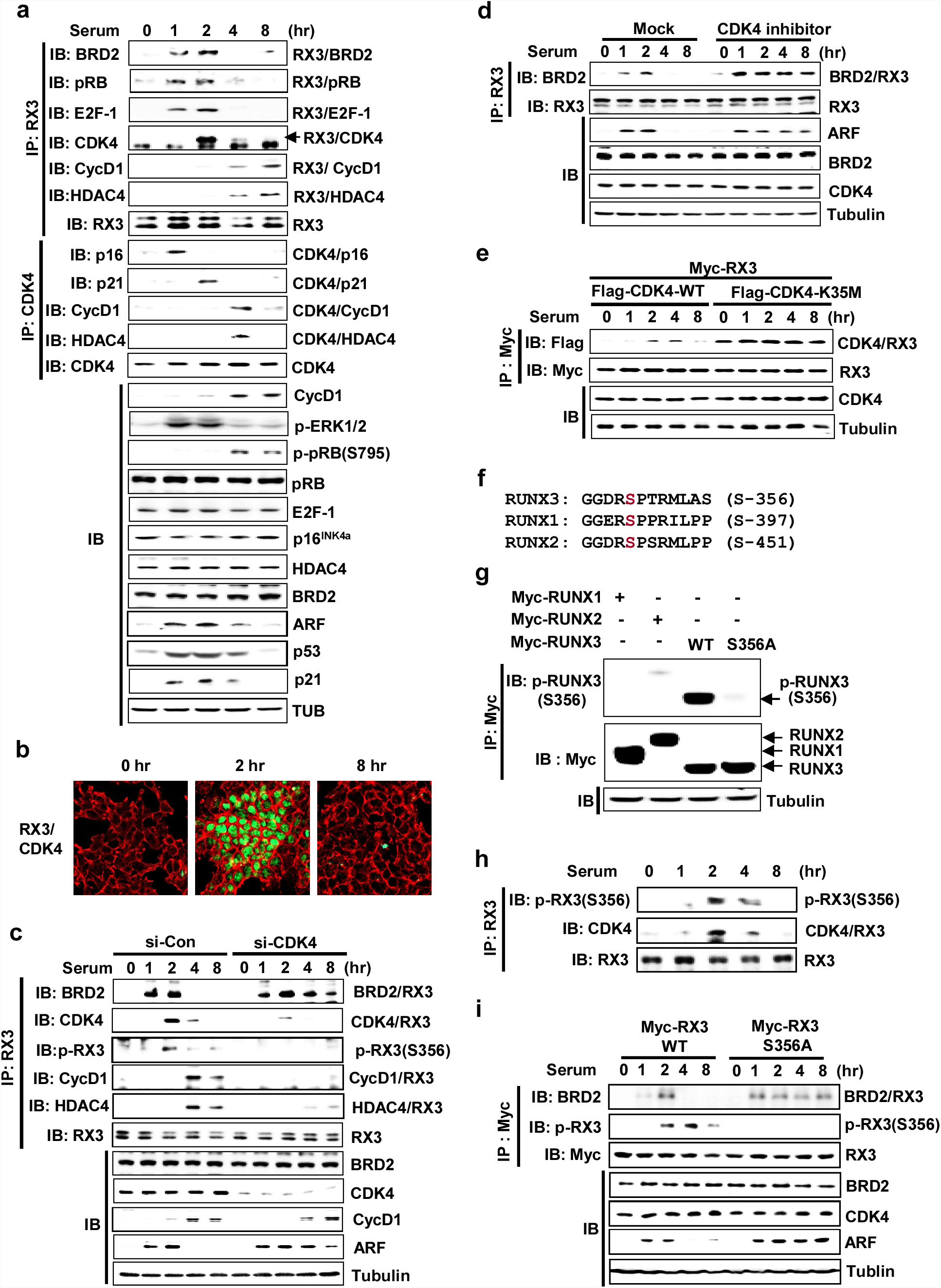
CDK4 plays key roles in the R-point transition. **a** HEK293 cells were serum-starved for 24 hr, stimulated with 10% serum, and harvested at the indicated time points. Time-dependent formation of the BRD2–RUNX3, pRB–RUNX3, E2F1–RUNX3, CDK4–RUNX3, Cyclin D1–RUNX3, HDAC4–RUNX3, p16^INK4a^–CDK4, p21–CDK4, Cyclin D1–CDK4, and HDAC4–CDK4 complexes was measured by IP and IB. RX3 formed a complex with BRD2, pRB, and E2F1 for 1–2 hr after serum stimulation. Time-dependent phosphorylation of pRB (at Ser-795) and ERK1/2 was measured by IB. **b** PLA assay showing the RUNX3–CDK interaction 2 hr after serum stimulation. **c** HEK293 cells were treated with control or CDK4-specific siRNA (si-con or si-CDK4), serum-starved for 24 hr, and then stimulated with serum for the indicated durations. Time-dependent formation of the BRD2–RUNX3, CDK4–RUNX3, HDAC4-RUNX3 and Cyclin D1–RUNX3 complexes and phosphorylated RUNX3 were measured by IP and IB. Time-dependent expression of ARF was measured by IB. **d** HEK293 cells were treated with CDK4 inhibitor (PD0332991, 500 nM), serum-starved for 24 hr, and then stimulated with serum for the indicated durations. Time-dependent formation of the BRD2–RUNX3 complex was measured by IP and IB. Time-dependent expression of ARF was measured by IB. **e** HEK293 cells were transfected with Myc-RUNX3 and Flag-CDK4 WT or Flag-CDK4-K35M (kinase inactive CDK4 mutant). Cells were serum-starved for 24 hr, stimulated with 10% serum, and harvested at the indicated time points. Time-dependent formation of the CDK4–RUNX3 complex was measured by IP and IB. **f** Comparison of peptide sequence adjacent to the CDK4-dependent phosphorylation site of RUNX family members. RUNX3 peptide phosphorylated at Ser-356 was used to raise rabbit anti-serum. **g** HEK293 cells were transfected with Myc-RUNX1, Myc-RUNX2, Myc-RUNX3 and Myc-RUNX3-S356A. At 2 hr after serum stimulation, cells were harvested, and RUNX3 phosphorylation at Ser-356 was analyzed by IP with anti-Myc antibody followed by IB with antiserum (anti-p-RUNX3-S356). **h** Cells were serum-starved for 24 hr, and then stimulated with 10% serum. Cells were harvested at the indicated time points. Time-dependent RUNX3–CDK4 interaction and RUNX3 phosphorylation at Ser-356 were measured by IP and IB. **i** HEK293 cells were transfected with Myc-RUNX3 or Myc-RUNX3-S356A, serum-starved for 24 hr, and then stimulated with 10% serum. Cells were harvested at the indicated time points. Time-dependent formation of the BRD2–RUNX3 complex, RUNX3 phosphorylation at Ser-356, and ARF expression were measured by IP and IB.

Although pRB and CDK4 were brought together within Rpa-RX3-AC 2 hr after serum stimulation, CDK4-mediated pRB phosphorylation occurred only 4 hr after stimulation (Fig. 4a). Analysis of the timing of interactions among CDK4 and its binding proteins revealed that CDK4 interacted with p16 1 hr after stimulation, and then replaced p16 with p21 at 2 hr (Fig. 4a). p21 facilitates the association of Cyclin D1 and CDK4 ^3,22,23^. However, Cyclin D1, which was induced 2 hr after stimulation, interacted with RNF2, but not with p21-bound CDK4 at that time point (Fig. 3a and Fig. 4a). The Cyclin D1–CDK4 interaction was detected after Rpa-RX3-TR formed (at 4 hr) (Fig. 4a). Notably, knockdown of *CDK4* markedly diminished the interactions of RUNX3 with Cyclin D1 and HDAC4 (Fig. 4c). Consequently, the RUNX3–BRD2 interaction and ARF expression were maintained for up to 8 hr (Fig. 4c). These results demonstrate that CDK4 of Rpa-RX3-AC and Cyclin D1 of PRC1-Cyclin D1–HDAC4 provide docking sites for the interaction of the two complexes, enabling formation of Rpa-RX3-TR.

### Cyclin D1–CDK4 triggers the R-point transition by phosphorylating RUNX3 as well as pRB

We then investigated the role of CDK4 kinase in R-point transition. Pharmacological inhibition of CDK4 activity maintained Rpa-RX3-AC (of which RUNX3–BRD2 is the core) and prolonged *ARF* expression for up to 8 hr (Fig. 4d). These results suggest that activation of CDK4 by the Cyclin D1–CDK4 interaction within Rpa-RX3-AC triggers the R-point transition.

Exogenously expressed Myc-RUNX3 and Flag-CDK4 also interacted 2–4 hr after serum stimulation, but dissociated thereafter (Fig. 4e). Interestingly, a kinase-dead CDK4 mutant (Flag-CDK4-K35M) bound to RUNX3 and did not dissociate for a long time after serum stimulation (Fig. 4e). These results demonstrate that CDK4 kinase activity is required for dissociation of CDK4 from RUNX3.

Parathyroid hormone-related protein decreases the level of RUNX3 in chondrocytes via CDK4-dependent RUNX3 phosphorylation at Ser-356 ^24^. Therefore, we investigated whether CDK4 also phosphorylates RUNX3 within Rpa-RX3-AC in order to promote the R-point transition. For this purpose, we raised rabbit polyclonal anti-serum against synthetic RUNX3 peptide phosphorylated at Ser-356 (Fig. 4f). Although the amino acid sequence of the RUNX3 peptide is similar to those of RUNX1 and RUNX2, the anti-serum specifically recognized only RUNX3 phosphorylated at Ser-356, but not RUNX1, RUNX2, or RUNX3-S356A (RUNX3 mutant lacking the CDK4-dependent phosphorylation site) (Fig. 4g). This anti-serum detected phosphorylation of endogenous RUNX3 contemporaneously with the RUNX3–CDK4 interaction (2–4 hr after serum stimulation) (Fig. 4h). RUNX3 phosphorylation was markedly diminished by knockdown of CDK4, implying that it is mediated by CDK4 (Fig. 4c). Notably, the interaction between exogenously expressed Myc-RUNX3-S356A and BRD2 and expression of ARF were maintained up to 8 hr after serum stimulation, whereas the interaction between Myc-RUNX3-WT and BRD2 was maintained only up to 2 hr afterward (Fig. 4i). These results demonstrate that CDK4 phosphorylates RUNX3, and that this modification triggers the release of BRD2 from RUNX3, which results in the R-point transition.

### Multiple signaling pathways downstream of RAS contribute to the R-point transition

The Cyclin D1–CDK4 interaction occurred 4 hr after serum stimulation, and Cyclin D1–CDK4-dependent pRB phosphorylation occurred at the same time point (Fig. 4a, CDK4-CycD1 and p-pRB (S-795)). Notably, however, CDK4-dependent RUNX3 phosphorylation occurred 2 hr after serum stimulation (Fig. 4c, h), earlier than the Cyclin D1-CDK4 interaction (Fig. 4a). These results suggest that CDK4 could be activated in a Cyclin D1-independent manner for RUNX3 phosphorylation. Activation of CDK4 depends upon CDK-activating kinase (CAK) activity ^25^. Recently, JNK was identified as one of the CAKs that phosphorylates CDK4 at T172 to activate it ^26^. Pharmacological inhibition of JNK activity markedly reduced RUNX3 phosphorylation at Ser-356 and maintained Rpa-RX3-AC (maintained interactions between RUNX3, CDK4, and BRD2) for up to 8 hr (Fig. 5a). Similarly, siRNA-mediated knockdown of JNK also decreased RUNX3 phosphorylation at Ser-356 and maintained Rpa-RX3-AC (Fig. 5b). In addition, exogenously expressed Flag-CDK4-WT phosphorylated Myc-RUNX3 2–4 hr after serum stimulation (Fig. 5c). However, Flag-CDK4-T172A (lacking the residue phosphorylated by JNK) failed to phosphorylate RUNX3 at Ser-356 (Fig. 5c). Flag-CDK4-T172A bound to RUNX3 even earlier, and did not dissociate for a long time after serum stimulation (Fig. 5c). Consistently, Flag-CDK4-T172A maintained Rpa-RX3-AC and prolonged ARF expression for up to 8 hr (Fig. 5c). These results demonstrate that the JNK pathway also contributes to the R-point transition by activating CDK4.

**Fig. 5:**
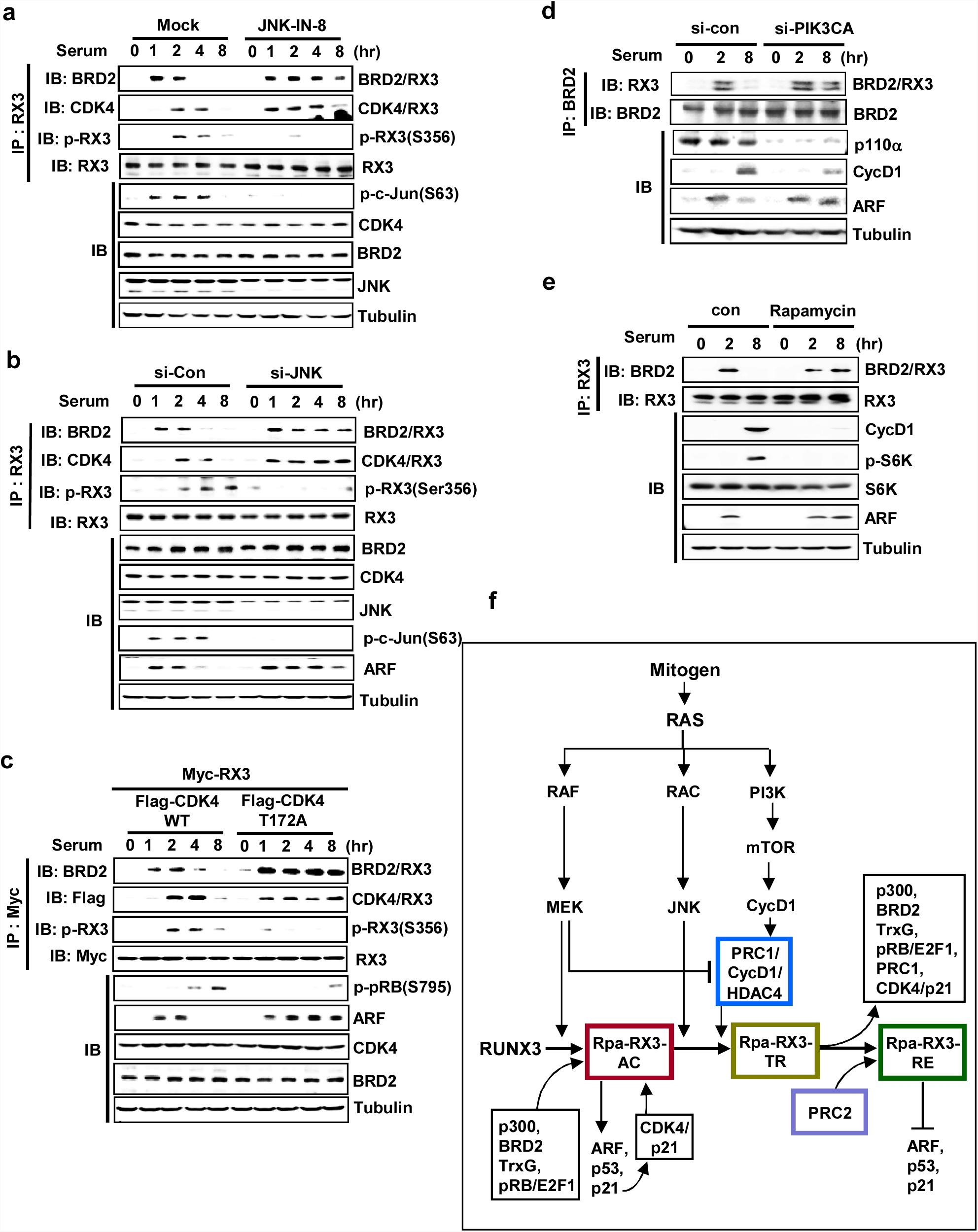
Multiple signals contribute to the R-point transition. **a** HEK293 cells were treated with JNK inhibitor (JNK-IN-8, 1 μM), serum-starved for 24 hr, and then stimulated with serum for the indicated durations. Time-dependent formation of the BRD2–RUNX3 and RUNX3–CDK4 complexes and phosphorylation of RUNX3 at Ser-356 were measured by IP and IB. Time-dependent expression of ARF was measured by IB. **b** HEK293 cells were treated with control or JNK-specific siRNA (si-con or si-JNK), serum-starved for 24 hr, and then stimulated with serum for the indicated durations. Time-dependent formation of the BRD2–RUNX3 and CDK4–RUNX3 complexes and phosphorylation of RUNX3 at Ser-356 were measured by IP and IB. Time-dependent expression of ARF was measured by IB. **c** HEK293 cells were transfected with Myc-RUNX3 and Flag-CDK4 WT or Flag-CDK4-T172A (CDK4 mutant defective in phosphorylation by JNK). Cells were serum-starved for 24 hr, and then stimulated with 10% serum. Time-dependent formation of the BRD2–RUNX3 and CDK4–RUNX3 complexes and phosphorylation of RUNX3 at Ser-356 were measured by IP and IB. Time-dependent phosphorylation of pRB and expression of ARF were measured by IB. **d** HEK293 cells were treated with control or PIK3CA-specific siRNA (si-con or si-PIK3CA), serum-starved for 24 hr, and then stimulated with serum for the indicated durations. Time-dependent formation of the BRD2–RUNX3 complex was measured by IP and IB. Time-dependent expression of ARF was measured by IB. **e** HEK293 cells were treated with control or mTORC1 inhibitor (Rapamycin, 100 nM), serum-starved for 24 hr, and then stimulated with serum for the indicated durations. Time-dependent formation of the BRD2–RUNX3 complex was measured by IP and IB. Time-dependent expression of ARF was measured by IB. Ribosomal protein S6 kinase beta-1 (S6K1), which is phosphorylated by mTOR signaling, was used for control. **f** Schematic illustration of the roles of the RAS signaling pathway in regulating the R-point transition. The RAS–RAF–MEK pathway facilitates formation of Rpa-RX3-AC and inhibits formation of the PRC1–Cyclin D1–HDAC4 complex. The RAS–RAC–JNK pathway activates CDK4. The RAS–PI3K pathway facilitates formation of Rpa-RX3-TR by contributing to translation of Cyclin D1.

Transcription and translation of Cyclin D1, which plays a key role in formation of Rpa-RX3-TR, are stimulated through the RAS–RAF and RAS–PI3K pathways, respectively ^3^. As expected, knockdown of the PI3K catalytic subunit (*PIK3CA*, encoding p110α) decreased the level of Cyclin D1 and maintained Rpa-RX3-AC (cored by RUNX3-BRD2) and prolonged ARF expression for up to 8 hr (Fig. 5d). Inhibition of mTOR, a downstream effecter of the RAS–PI3K pathway, by Rapamycin also maintained Rpa-RX3-AC (cored by RUNX3-BRD2) and prolonged ARF expression for up to 8 hr (Fig. 5e). These results demonstrate that the PI3K pathway also contributes to the transition from Rpa-RX3-AC to Rpa-RX3-TR by inducing Cyclin D1.

We previously showed that RAS-RAF-MEK pathway triggers RUNX3-BRD2 interaction ^14^. Therefore, the three major pathways downstream of RAS (MEK, JNK, and PI3K) contribute to the R-point transition at distinct stages. These results demonstrate that cells cannot pass through the R-point and progress toward S phase if RAS or its downstream signals are deregulated. This molecular program suggests that the R-point decision includes an oncogene surveillance mechanism. The contributions of these pathways to each stage of the R-point are summarized in Fig. 5f. An overview of the R-point transition and R-point–associated chromatin dynamics in response to normal mitogenic signals and oncogenic RAS signals is provided in Supplementary Fig. 4.

### Oncogenic K-RAS inhibits the R-point transition by inhibiting Rpa-RX3-TR formation

Our results show that the transition from Rpa-RX3-AC to Rpa-RX3-TR occurs only after MEK activity is downregulated (Fig. 4a, p-ERK1/2). Therefore, we asked what would happen if the RAS-RAF-MEK pathway was constitutively activated. Ectopic expression of oncogenic *K-RAS* (Myc-K-RAS^G12V^) facilitated the interactions of RUNX3 with p300, BRD2, SWI/SNF, TFIID, and CDK4, and maintained the complex for up to 8 hr (Fig. 6a). By contrast, expression of Myc-K-RAS^G12V^ inhibited the interactions between RUNX3, Cyclin D1, and HDAC4, as well as the interaction between BRD2 and PRC1 (Fig. 6a). ChIP analysis also showed that expression of oncogenic K-RAS maintained binding of Rpa-RX3-AC to the *ARF* locus, but inhibited binding of Cyclin D1, HDAC4 and PRC1 (Fig. 6b). These results suggest that oncogenic *K-RAS* facilitates Rpa-RX3-AC formation but inhibits Rpa-RX3-TR formation. In addition, oncogenic *K-RAS* facilitated formation of the PRC1–Cyclin D1 complex (Fig. 6a, RNF2–CycD1), but inhibited incorporation of HDAC4 into the complex (Fig. 6a, HDAC4–CycD1). Therefore, oncogenic *K-RAS* inhibits Rpa-RX3-TR formation by inhibiting assembly of HDAC4 into the PRC1–Cyclin D1 complex.

**Fig. 6:**
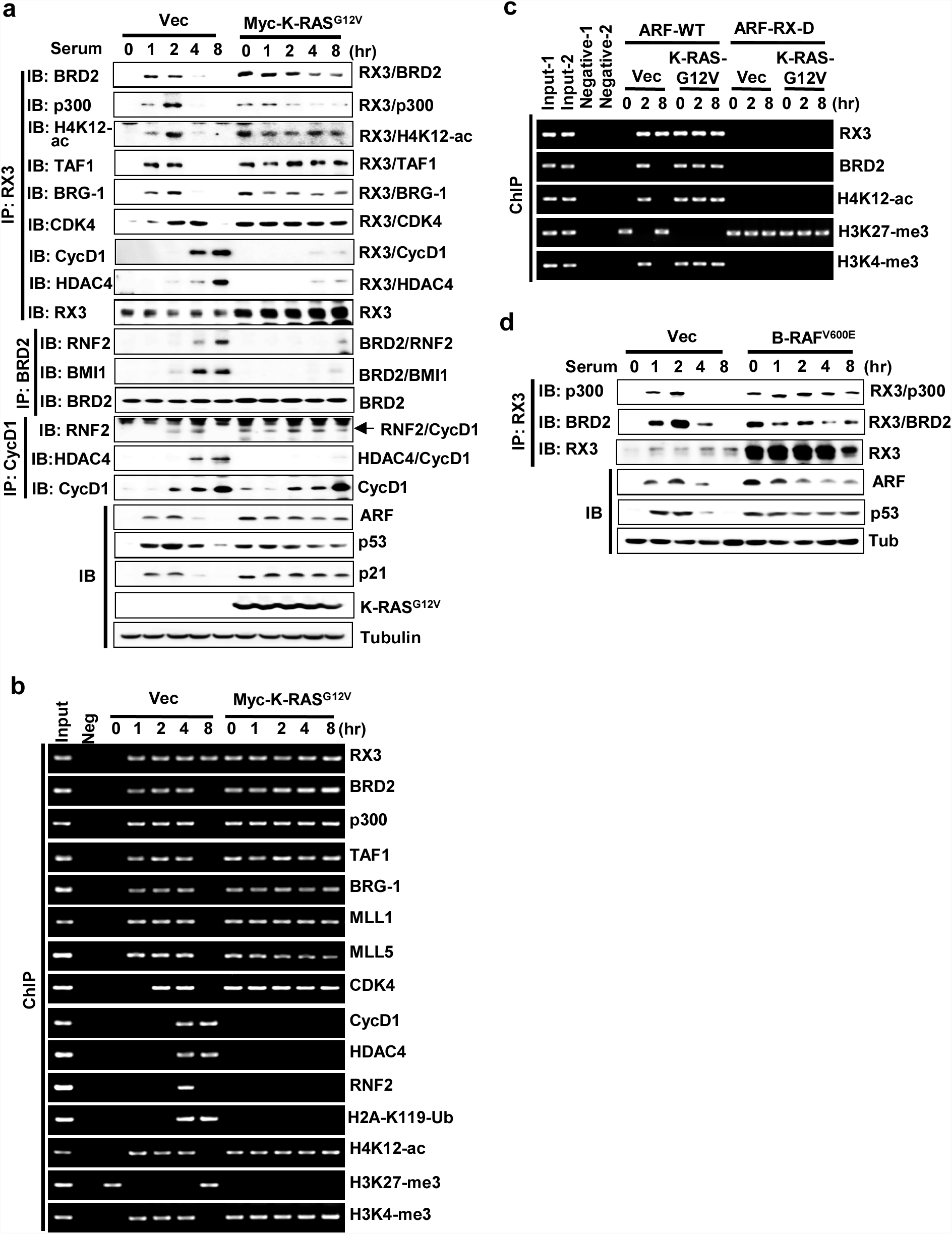
R-point surveils aberrant oncogene activation through the Rpa-transition program. **a** HEK293 cells were transfected with empty vector (Vec) or Myc-K-RAS^G12V^, serum-starved for 24 hr, stimulated with 10% serum, and harvested at the indicated time points. The time-dependent interactions among the components of Rpa-RX3-AC, Rpa-RX3-TR, and Rpa-RX3-RE were measured by IP and IB. Expression levels of ARF, p53, p21, and Myc-K-Ras^G12V^ were measured by IB. **b** HEK293 cells were serum-starved for 24 hr, and then stimulated with 10% serum. Binding of the components of Rpa-RX3-AC, Rpa-RX3-TR, and Rpa-RX3-RE to the *p14^ARF^* promoter and histone marks (H4K12-ac, H3K27-me3, H3K4-me3, and H2A-K119-Ubi) at the locus were measured by ChIP at the indicated time points. One-thirtieth of the lysates were PCR-amplified as input samples. **c** Wild-type HEK293 cells (HEK293-ARF-WT) and HEK293-ARF-RX-D cells were transfected with empty vector (Vec) or Myc-K-Ras^G12V^, serum-starved for 24 hr, and then stimulated with 10% serum. Cells were harvested at the indicated time points, and the binding of RUNX3, BRD2, H4K12-ac, H3K27-me3, and H3K4-me3 to the *ARF* promoter was measured by ChIP at the indicated time points. One-thirtieth of the lysates were PCR-amplified as input samples. **d** HEK293 cells were transfected with empty vector (Vec) or B-RAF^V600E^, serum-starved for 24 hr, stimulated with 10% serum, and harvested at the indicated time points. Time-dependent formation of the RUNX3–p300 and BRD2–RUNX3 complexes was measured by IP and IB. Time-dependent expression of ARF and p53 was measured by IB.

ChIP analysis revealed that H4K12-ac and H3K4-me3 histone marks were maintained for a long time by expression of Myc-K-RAS^G12V^ (Fig. 6b). Consistently, when Myc-K-RAS^G12V^ was expressed, ARF and p21 (targets of Rpa-RX3-AC) were not downregulated, but were instead maintained for long periods (Fig. 6a). Prolonged binding of Rpa-RX3-AC and maintenance of H4K12-ac and H3K4-me3 histone marks at the *ARF* locus by expression of oncogenic *K-RAS* were abolished by deletion of the RUNX-binding site from the locus (Fig. 6c).

In addition to oncogenic *K-RAS*, oncogenic *B-Raf* (B-RAF^V600E^) also maintained Rpa-RX3-AC, prolonged expression of ARF, and stabilized p53 for long periods (Fig. 6d). These results demonstrate that the *ARF* locus is opened by Rpa-RX3-AC and closed at the R-point in normal cells, but is not closed in cells expressing oncogenic *K-RAS* or oncogenic *B-RAF* due to failure of the Rpa-RX3-AC → Rpa-RX3-TR transition.

### *RUNX3* suppresses endogenous oncogenic *K-RAS*–driven oncogenesis through R-point-associated programs

We next investigated the protective role of RUNX3 against endogenous oncogenic *K-RAS* in H460 human lung cancer cells (*K-RAS*–activated without amplification, *RUNX3*-inactivated, *ARF* wild-type, and p53 wild-type). By stable transfection, we obtained cell lines expressing H460-vec, H460-ERT2-RUNX3 (expressing ERT2 fused with wild-type RUNX3), and H460-ERT2-RUNX3-K94/171R [expressing ERT2 fused with RUNX3 mutant, which does not interact with BRD2 ^14^]. The ERT2 fusion proteins remained in the cytoplasm until 4 hr after induction by 4-hydroxytamoxifen (4-OHT), but were localized to the nucleus by 8 hr (Fig. 7a). In the absence of inducer, none of the cell lines formed Rpa-RX3-AC at any time point after serum stimulation (Fig. 7b), even though they expressed oncogenic *K-RAS*. At 8 hr after inducer treatment, H460-ERT2-RUNX3 cells, but not H460-ERT2-RUNX3-K94/171R cells, formed Rpa-RX3-AC (Fig. 7b). In inducer-treated H460-ERT2-RUNX3 cells, Rpa-RX3-AC was maintained and the ARF-p53 pathway was activated for a long time because oncogenic *K-RAS* maintained Rpa-RX3-AC (Fig. 7b). ChIP analysis confirmed that Rpa-RX3-AC bound to its target locus for a long time in inducer-treated H460-ERT2-RUNX3 cells (Fig. 7c). Consistently, H4K12-ac and H3K4-me3 were enriched, whereas H3K27-me3 was diminished, in H460-ERT2-RUNX3 cells, but not in H460-ERT2-RUNX3-K94/171R cells, 8–16 hr after induction (Fig. 7c). These results indicate that inducer-treated H460-ERT2-RUNX3 cells clearly responded to endogenous oncogenic *K-RAS* via the R-point defense program, but this process was not initiated in H460-ERT2-RUNX3-K94/171R cells and parental H460 cells, which lack *RUNX3* expression.

**Fig. 7:**
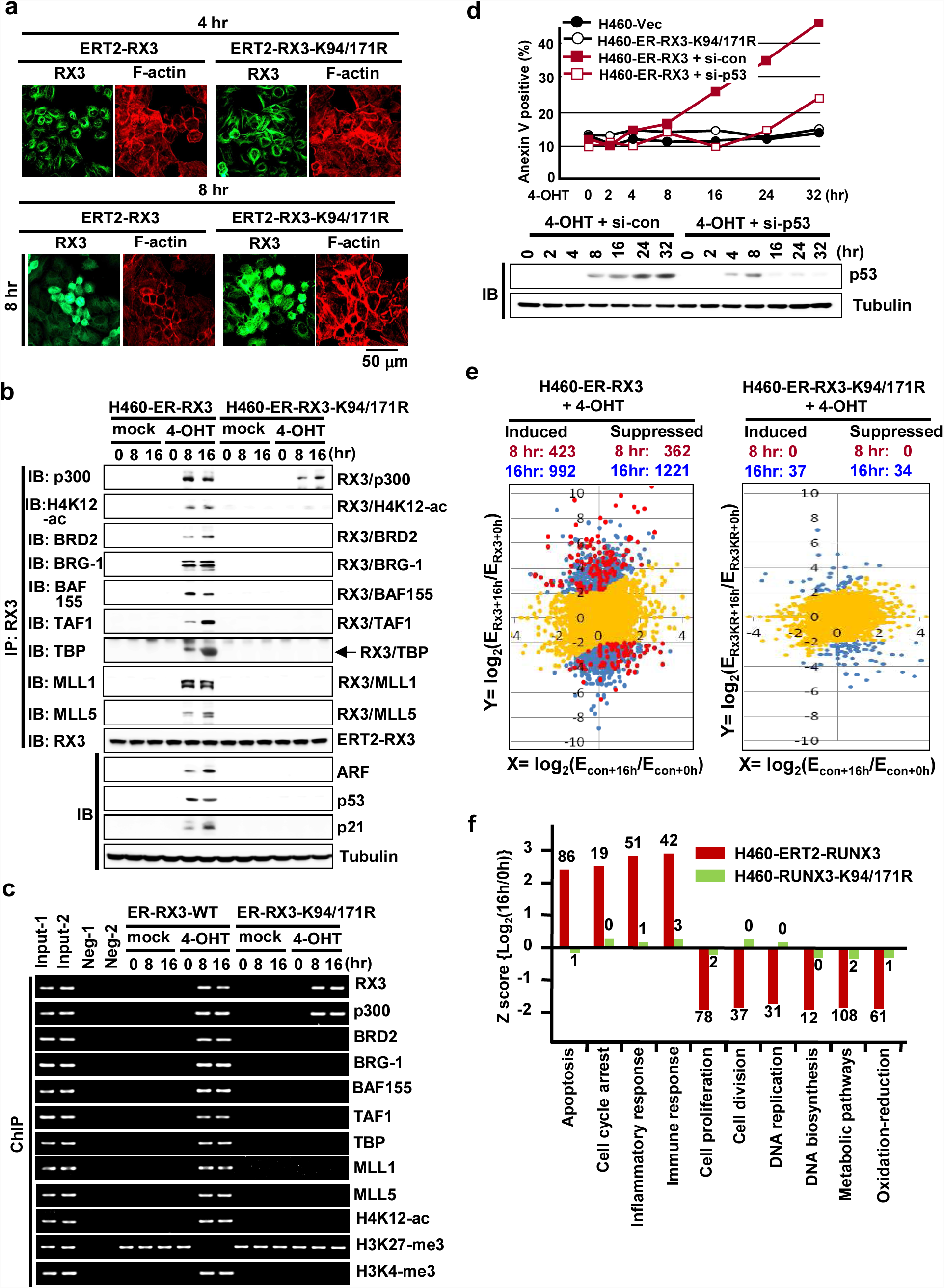
RUNX3 defends against endogenous oncogenic *K-Ras* by contributing to the R-point decision. **a** H460-ERT2-RUNX3 and H460-ERT2-RUNX3-K94/171R cells were synchronized by serum deprivation and stimulated with 10% serum and 1 μM 4-OHT for the indicated durations (0, 4, and 8 hr). Time-dependent subcellular localization of the expressed proteins was analyzed by double immunofluorescence staining (green=RUNX3; red=F-actin). **b-c** H460-ERT2-RUNX3 and H460-ERT2-RUNX3-K94/171R cells were serum-starved for 24 hr, stimulated with 10% serum or 10% serum + 1 μM 4-OHT. Cells were harvested at the indicated time points, and the time-dependent interactions of RUNX3 with BRD2, p300, H4K12-ac, TFIID complex (TAF1, TAF7, and TBP), SWI/SNF complex (BRG-1 and BAF155), and MLL1/5 were measured by IP and IB. Expression of p14^ARF^, p53, and p21 was measured by IB. The binding of the proteins and H4K12-ac, H3K27-me3, and H3K4-me3 to the *ARF* promoter was measured by ChIP at the indicated time points. One-thirtieth of the lysates were PCR-amplified as input samples. **d** H460-vec, H460-ERT2-RUNX3, and H460-ERT2-RUNX3-K94/171R cells were treated with indicated si-RNA, serum-starved for 24 hr, and then stimulated with 10% serum or 10% serum + 1 μM 4-OHT for the indicated durations. Apoptotic cells were detected by flow cytometry after Annexin V–FITC/PI staining. The levels of p53 were measured by IB. **e** H460-vec, H460-ERT2-RUNX3, and H460-ERT2-RUNX3-K94/171R cells were serum-starved for 24 hr, and then stimulated with 10% serum or 10% serum + 1 μM 4-OHT for 0, 8, or 16 hr. RNA was extracted from the cells, and gene expression patterns were analyzed by mRNA sequencing. Expression of genes 8 or 16 hr after serum stimulation was quantified as log_2_(fold change) relative to the average of control reactions (i.e., before serum stimulation, 0 hr) for each cell line. Differential gene expression in response to expression of wild-type RUNX3 was analyzed by plotting log_2_(E_Rx3+16h_/E_Rx3+0h_) and log_2_(E_vec+16h_/E_vec+0h_); the results are shown on the left. Differential gene expression changes in response to expression of RUNX3-K94/171R were analyzed by plotting log_2_(E_Rx3KR+16h_/E_Rx3KR+0h_) and log_2_(E_vec+16h_/E_vec+0h_); the results are shown on the right. E_Rx3+16h_ and E_Rx3+0h_, E_Rx3KR+16h_ and E_Rx3KR+0h_, and E_vec-16h_ and E_vec-0h_ are the average expression levels of genes 0 or 16 hr after 4-OHT stimulation in H460-ERT2-RUNX3, H460-ERT2-RUNX3-K94/171R, and H460-vec cells, respectively. Yellow spots indicate genes regulated in a *RUNX3*-independent manner. Red and blue spots indicate genes regulated in a *RUNX3*-dependent manner 8 and 16 hr after 4-OHT stimulation, respectively (FDR < 0.001, p < 0.05). **f** RUNX3-dependent genes involved in various signaling pathways were analyzed using the DAVID Bioinformatics Resources 6.8 ^34^. Gene expression levels 16 hr after 4-OHT stimulation in the indicated cells were quantified as the fold change relative to the average of un-stimulated levels. Z-scores of log_2_(E_Rx3+16h_/E_Rx3+0h_) and log_2_(E_Rx3KR+16h_/E_Rx3KR+0h_) for major signaling pathways are shown for H460-ERT2-RUNX3 and H460-ERT2-RUNX3-K94/171R cells. Numbers at the top or bottom of the bars indicate the numbers of genes significantly up- or downregulated by inducer treatment. All categories were enriched with p < 0.05.

Consequently, H460-ERT2-RUNX3 cells, but not H460-ERT2-RUNX3-K94/171R cells, underwent apoptosis after inducer treatment (Video 1). FACS analysis at various time points after inducer treatment also revealed that the apoptosis rate of H460-ERT2-RUNX3 cells began to increase 16 hr after inducer treatment, and increased further thereafter (Fig. 7d). Apoptosis was markedly suppressed by siRNA-mediated knockdown of p53 (Fig. 7d and Supplementary Fig. 5). H460-vec cells and H460-ERT2-RUNX3-K94/171R cells did not undergo apoptosis regardless of the presence or absence of inducer (Fig. 7d). These results demonstrate that Rpa-RX3-AC defends the cell against endogenous oncogenic *K-RAS*, and that the ARF–p53 pathway is engaged in the R-point defense program.

### The R-point governs multiple programs of tumor suppression

To identify genes regulated by Rpa-RX3-AC at the R-point, we performed mRNA sequencing (RNA-seq) in H460-ERT2-RUNX3 and H460-ERT2-RUNX3-K94/171R cells. H460-vec cells were used as controls to eliminate any effect of inducer alone. Two-dimensional plots of expression changes in response to inducer treatment revealed that 423 and 362 genes were induced and suppressed, respectively [|log_2_(fold change)| ≥ 2], in H460-ERT2-RUNX3 cells 8 hr after inducer treatment (Fig. 7e, red spots). The numbers of genes induced and suppressed increased to 992 and 1221, respectively, after 16 hr (Fig. 7e, blue spots). By contrast, very few genes were affected by inducer treatment in H460-ERT2-RUNX3-K94/171R cells (Fig. 7e). These results suggest that Rpa-RX3-AC mediated most of the up- and downregulation of genes in inducer-treated H460-ERT2-RUNX3 cells. Because RUNX3 does not form the repressor complex (Rpa-RX3-RE) in H460 cells due to the oncogenic *K-RAS* mutation, the repressed genes were likely indirect targets of RUNX3. These genes might have been inappropriately activated in H460 cells due to disruption of the R-point, but then suppressed by restoration of the R-point after *RUNX3* expression.

Analysis of the Z-scores of major RUNX3-dependent signaling pathways revealed that genes involved in apoptosis, cell-cycle arrest, inflammatory response, and immune response were induced (Fig. 7f). On the other hand, genes involved in cell proliferation, DNA replication, and metabolic pathways were suppressed by RUNX3 expression (Fig. 7f). These results suggest that the R-point defends against oncogenic *K-RAS*–induced tumorigenesis not only by regulating intracellular programs (cell cycle, apoptosis, and metabolic pathways), but also by regulating extracellular programs (inflammatory response and immune response). *RUNX3*-dependent up-and downregulated genes involved in major signaling pathways are listed in Supplementary Fig. 6. Detailed RNA-seq results are provided in the Excel file H460-RUNX3.xlsx (Supplementary information).

## Discussion

Appropriate decision-making at the R-point, which is critical for tumor suppression, is achieved by transcriptional regulation of a large number of R-point– associated genes. The molecular mechanism by which cells regulate R-point–associated gene expression to make a signal-dependent R-point decision is one of the most fundamental issues in cell biology. In this study, we showed that RUNX3 transactivates R-point–associated genes by binding to its target loci and opening chromatin structure by sequential recruitment of mediator, chromatin-remodeling complex, basal transcription complex, histone modifiers, and cell-cycle regulators, which together form Rpa-RX3-AC. Briefly, between 1 and 2 hr after serum stimulation, RUNX3 binds to target loci and recruits various proteins including p300, E2F1, pRB, BRD2, H4K12-ac, SWI/SNF, TFIID, MLL1/5, and CDK4 to form Rpa-RX3-AC, which opens the chromatin. Cyclin D1, which is induced 2 hr after serum stimulation, interacts with PRC1 (containing RNF2) and matures into the PRC1–CyclinD1–HDAC4 complex. Between 2 and 4 hr after serum stimulation, the Rpa-RX3-AC and PRC1–CyclinD1– HDAC4 complexes interact to form a transient complex, Rpa-RX3-TR. Subsequently, Rpa-RX3-TR is destroyed (at 4 hr) to form Rpa-RX3-RE and PRC1-BRD2 (at 8 hr). Rpa-RX3-RE, containing RUNX3, Cyclin D1, HDAC4, and PRC2 (containing EZH2), continues to bind target loci in order to suppress gene expression, whereas PRC1–BRD2 is released from chromatin. Therefore, our results identify RUNX3 as a pioneer factor of the R-point and reveal the molecular mechanisms by which appropriate chromatin remodelers and histone modifiers are selectively recruited to target loci in response to mitogenic signals.

The function of pioneer factors in regulation of chromatin structure was originally described in the context of development. Orchestration of gene expression to direct embryonic development includes the activities of PcG proteins within the PRC1 and PRC2 complexes, as well as TrxG proteins within COMPASS (complex of proteins associated with Set1) ^27^. Because embryonic development is a long-term process, most studies to date have been based on static biochemical investigations. By contrast, R-point–associated genes are dynamically regulated by R-point–associated complexes (Rpa-RX3-AC, Rpa-RX3-TR, and Rpa-RX3-RE) during each cell cycle. Although both Rpa-RX3-AC and COMPASS contain TrxG proteins, these complexes are distinct. The major differences are that (1) Rpa-RX3-AC is assembled only at target chromatin loci, Rpa-RX3-AC interacts with PcG proteins to form Rpa-RX3-TR at the R-point, and assembly/disassembly of Rpa-RX3-AC is controlled by cell-cycle regulators, whereas COMPASS is not. Therefore, identification of RUNX3, a pioneer factor of the R-point, as the core of Rpa-RX3-AC reveals a new mechanism underlying the dynamic regulation of R-point–associated genes.

The emerging picture of chromatin function in cancer involves complex interplay of chromatin-modifying enzymes. In general, loss of TrxG and gain of PcG is a common theme in human cancer, demonstrating the respective tumor-suppressive and oncogenic roles of these proteins ^28^. This is also consistent with our claim that the R-point constitutes an oncogene surveillance mechanism: TrxG could suppress tumors through R-point regulation as a component of Rpa-RX3-AC, and PcG could drive cell cycle progression by destroying Rpa-RX3-AC.

Although TrxG and PcG play opposing roles in transcriptional programs ^28^, some promoters are associated with both marks (H3K4-me3 and H3K27-me3), a phenomenon known as bivalent modification ^29–31^. For example, *ARF* is a bivalent gene ^32^. It is worth mentioning that most bivalent genes were identified by population analysis in unsynchronized cells. Our results show that the chromatin structure of the *ARF* locus is dynamically regulated during the cell cycle. Therefore, the chromatin structure of some bivalent genes may be dynamically regulated during the cell cycle.

Our results show that the R-point decision-making program is tightly regulated by the concerted action of multiple pathways that mediate mitogenic signaling. If the balance of mitogenic signals is deregulated, the cell cannot pass through the R-point. Therefore, the R-point constitutes an oncogene surveillance mechanism that can discriminate between normal mitogenic and abnormal oncogenic signals. It is widely accepted that the ARF–p53 pathway plays major roles in oncogene surveillance mechanism. We previously showed that *Runx3* is downregulated in most of *K-Ras*– activated human and mouse lung ADC cells ^14^. The H460 human lung cancer cell line represents such a case (*K-RAS*–activated, *RUNX3*-inactivated, *ARF* wild-type, and p53 wild-type). Using this cell line, we showed that RUNX3 plays a central role in triggering the ARF–p53 pathway in response to oncogene activation by forming Rpa-RX3-AC, a key regulator of the R-point. Therefore, our results explain that some *K-RAS*–activated lung ADC cells can proliferate in the absence of RUNX3 because the p53 pathway is not activated in the absence of the R-point. Thus, our results demonstrate that the R-point constitutes an oncogene surveillance mechanism and explain why the R-point is perturbed in nearly all cancer cells. It is worth emphasizing that in multiple kinds of tumors, *RUNX3* is frequently inactivated by epigenetic alterations ^13^, which could in theory be reversed ^33^. Therefore, RUNX3 represents a therapeutic target for multiple types of tumors.

## ACKNOWLEDGMENTS

S-C Bae is supported by a Creative Research Grant (2014R1A3A2030690) through the National Research Foundation (NRF) of Korea. J-W Lee is supported by Basic Science Research Program grant 2018R1C1B6001532, Y-S Lee is supported by Basic Science Research Program grant 2017R1D1A3B03034076, J-W Jang is supported by Basic Science Research Program grant 2018R1C1B6009179, and J-W Hyun is supported by Basic Research Laboratory Program grant NRF-2017R1A4A1014512 through the NRF, funded by the Ministry of Science, ICT and future Planning (MSIP) of Korea government. D-M Kim was supported by Korean Foundation of Cancer Research (KFCR-2016-003). I-Y Park was supported by “Cooperative Research Program for Agriculture Science and Technology Development (PJ01226101)” Rural Development Administration, Republic of Korea. Y Ito is supported by the National Research Foundation Singapore and the Singapore Ministry of Education under its Research Centres of Excellence initiative

## AUTHOR CONTRIBUTIONS

J-W Lee and D-M Kim analyzed assembly and disassembly of the R-point–associated complexes. J-W Jang analyzed fly genetics. Y-S Lee, X-Z Chi, I-Y Park and J-W Hyun performed yeast two-hybrid screening. T-G Park performed the 3C assay. Y Ito and S-C Bae interpreted the results and wrote the manuscript. All authors contributed to the editing of the manuscript.

## COMPETING FINANCIAL INTERESTS

The authors declare no competing interests.

## SUPPLEMENTAL INFORMATION

Supplemental information includes 6 Supplementary Figures and an Excel file.

## CONTACT FOR REAGENT AND RESOURCE SHARING

Further information and requests for resources and reagents should be directed to and will be fulfilled by the Lead Contact, Suk-Chul Bae.

## Methods

### Cell lines

HEK293 cells were maintained in Dulbecco’s modified Eagle’s medium (Gibco BRL, Thermo Fisher Scientific, Waltham, MA, USA) supplemented with 10% fetal bovine serum (Gibco BRL) and 1% penicillin/streptomycin (Invitrogen, Carlsbad, CA, USA). H460 cells and H460 stable cell lines were maintained in RPMI 1640 medium (Gibco BRL) supplemented with 10% fetal bovine serum (Gibco BRL) and 1% penicillin/streptomycin (Invitrogen). All cell lines were incubated at 37°C with 5% CO_2_.

### *Drosophila* genetics

*GMR-Gal4* was obtained from the Bloomington Drosophila Stock Center (Bloomington, IN, USA). *UAS-L*z was kindly provided by U. Banerjee (University of California, Los Angeles, CA, USA) ^35^. An EP line harboring an enhancer P element insertion within the upstream regulatory region of *Trithorax* (G14137) was obtained from GeneExel (KIST, Daejeon, South Korea). *Drosophila* stocks were maintained and cultured on standard cornmeal–yeast–agar medium at 25°C.

### DNA transfection, IP, and IB

Transient transfections in all cell lines were performed using Lipofectamine Plus reagent and Lipofectamine (Invitrogen). Cell lysates were incubated with the appropriate monoclonal antibodies for 3 hr at 4°C, and then with protein G–Sepharose beads (Amersham Pharmacia Biotech, Piscataway, NJ, USA) for 1 hr at 4°C. For detection of endogenous proteins, lysates were incubated with the appropriate monoclonal antibodies for 6–12 hr at 4°C, and then with protein G–Sepharose beads (Amersham Pharmacia Biotech) for 3 hr at 4°C. Immunoprecipitates were resolved on SDS–polyacrylamide gel electrophoresis (SDS-PAGE) gels and transferred to a polyvinylidene difluoride membrane (Millipore, Billerica, MA, USA). The membrane was immunoblotted with the appropriate antibodies after blocking and visualized on an Amersham™ Imager 600 (GE Healthcare, Chicago, IL, USA) after treatment with ECL solution (Amersham Pharmacia Biotech).

### Antibodies

Antibodies targeting Cyclin D1, CDK4, HDAC4, p300, p53, p21, p14, E2F1, TAF1, TAF7, TBP, BRG-1, and BAF155 were obtained from Santa Cruz Biotechnology (Dallas, TX, USA). Antibodies targeting H2AK119-u1, H3K412-ac, H3K27-me3, H3K4-me3, RNF2, BMI1, EZH2, phospho-pRB (Ser-795), phospho-ERK1/2, and Acetylated Lys were obtained from Cell Signaling Technology (Danvers, MA, USA). Antibodies targeting RUNX3 (5G4), p16 and EED were obtained from Abcam (Cambridge, UK). Antibodies targeting HA (12CA5; Roche Applied Science, Mannheim, Germany), FLAG (M2; Sigma, MO, USA), Myc (9E10; Santa Cruz Biotechnology), BRD2 (M01; Abnova, Taipei City, Taiwan), pRB (BD Biosciences, Seoul, South Korea), MLL5 (St John’s Laboratory, London, UK) and MLL1 (Bethyl Laboratories Inc., TX, USA) were used for IB and IP. Anti RUNX3-phospho-S356 was made rabbit polyclonal anti-serum against synthetic RUNX3 peptide phosphorylated at Ser-356.

### *In vitro* translation

The TnT^®^ Coupled Reticulocyte Lysate System (Promega, Madison, WI, USA) is available in two configurations, for transcription and translation of genes cloned downstream of either the T7 or SP6 RNA polymerase promoters. To use these systems, 2.0 μg of circular plasmid DNA containing a SP6 promoter was added directly to TnT^®^ lysate and TnT^®^ Quick Master Mix, and then incubated in a 50 μl reaction volume for 1.5 hr at 30°C. Synthesized proteins were analyzed by SDS-PAGE.

### Proximity ligation assay (PLA)

The PLA was performed using the Duolink^®^ In Situ PLA^®^ Kit (Sigma, St. Louis, MO, USA). Briefly, cells were grown, fixed, and permeabilized. The samples were then incubated overnight at 4°C with primary antibodies against the two proteins to be examined, washed [Buffer A: 0.01 M Tris-HCl (pH 7.4), 0.15 M NaCl, and 0.05% Tween 20], incubated at 37°C for 60 min with specific probes, stained for F-actin to visualize cytoplasm, and washed with Buffer B [0.2 M Tris-HCl (pH 7.5) and 0.1 M NaCl]. Signals were visualized as distinct fluorescent spots on a fluorescence microscope (Carl Zeiss AXIO Zoom.V16 and ApoTome.2). Background correction, contrast adjustment of raw images, and quantification of fluorescence signals were performed using the Zen 2012 Blue Edition software (Carl Zeiss, Oberkochen, Germany).

### Inhibitor and siRNA

CDK4 inhibitor (PD0332991), JNK inhibitor (JNK-IN-8) and Rapamycin (R8781) were purchased from Sigma-Aldrich. Cells were treated with CDK4 inhibitor (500 nM), JNK inhibitor (1 μM) or Rapamycin (100 nM), and harvested at the indicated time points after serum stimulation. Knockdown analysis was performed by transfecting HEK293 cells with 50 nM siRNA using RNAiMAX (Invitrogen, CA, USA) before serum starvation. Cells were harvested at the indicated time points after serum stimulation. BRD2, p53, CDK4, RNF2 and Cyclin D1 siRNAs were purchased from Bioneer (Daejeon, South Korea). HDAC4 siRNAs was purchased from Cell Signaling Technology. Sequences of siRNAs were as follows:

(si-BRD2 sense: CACUUGGCCUGCAUGACUA

antisense: UAGUCAUGCAGGCCAAGUG)

(si-p53 sense: CAGUUUGAGGUGCGUGUU

antisense: AACACGCACCUCAAAGCUG)

(si-CDK4 sense: CCAGAAUCUACAGCUACCA

antisense: UGGUAGCUGUAGAUUCCUGG)

(si-Cyclin D1 sense: GACCUUCGUUGCCCUCUGU

antisense: ACAGAGGGCAACGAAGGUC)

(si-RNF2 sense: UGAUAGGGUAUUGAGUGUA

antisense: UACACUCAAUACCCUAUCA)

### Deletion of the RUNX-binding site in the *CDKN2A* (*p14^ARF^*) enhancer region

To delete the RUNX-binding site (GACCGCA) in the *CDKN2A* (*p14^ARF^*) enhancer region (ntd −1466) by the CRISPR/Cas9 method, HEK293 cells were transfected with the pRGEN-CDKN2A target plasmid (target sequence: 5’-GACGGATCCAGGCAGACCGCAGG) and pRGEN-Cas9-Hyg-CMV (ToolGen, Seoul, South Korea). The cells were maintained in standard culture medium (10% Dulbecco’s modified Eagle’s medium) containing 800 μg/ml hygromycin B (H3274; Sigma). Hygromycin B-resistant cells were selected. Deletion of the RUNX3-binding site was confirmed by nucleotide sequencing.

### ChIP assay

ChIP assays were performed using the ChIP assay kit (cat # 17-295; Millipore). HEK293 cells or H460-derived stable cell lines were serum-starved for 24 hr, treated with 10% serum or 10% serum/1 μM 4-OHT (Sigma), harvested at the indicated time points, and cross-linked with formaldehyde (1% [v/v,] 10 min, 37°C). Chromatin was immunoprecipitated with the indicated antibodies. The *p14* promoter region was amplified by PCR using the following primers (previously reported primers) ^20^. (*p14^ARF^*-Forward: AGTGGCTACGTAAGAGTGATCGC) (*p14^ARF^*-Reverse: CTTACAGATCAGACGTCAAGCCC)

### Chromosome Conformation Capture assay (3C assay)

3C assays were performed as described previously ^21^. Briefly, 1.0 × 10^7^ cells were cross-linked using 2% formaldehyde in 10 ml of PBS containing 10% fetal bovine serum and incubated for 10 min at room temperature. The reaction tubes were quenched with 1.425 ml of 1 M glycine. The fixed cells were washed twice with ice-cold PBS and then harvested using 1 ml of ice-cold PBS. Harvested cells were re-suspended in 5 ml of cold lysis buffer [10 mM Tris-HCl (pH 7.5), 10 mM NaCl, 0.2% NP-40, and protease inhibitors] and incubated for 10 min on ice. The lysates were centrifuged at 400 *g* for 10 min at 4°C. The pelleted nuclei were washed with 0.5 ml of 1.2× restriction enzyme buffer, and then re-suspended in 0.5 ml of the same buffer. SDS was added to a final concentration of 0.3%, and pelleted nuclei were incubated for 1 hr at 37°C. Triton X-100 was added to a final concentration of 2% to quench the SDS, and the nuclei were incubated for 1 hr at 37°C. Next, 400 U of *Xba*I was added, and the sample was incubated overnight at 37°C. SDS was added to a concentration of 1.6%, and the sample was incubated for 25 min at 65°C. Digested genomic DNA was suspended in 6.125 ml of ice-cold 1.15× ligation buffer containing 1% Triton X-100. T4 DNA ligase (100 U) was added, and the reaction mixtures were incubated for 4 hr at 16°C, followed by 30 min at room temperature. Reaction mixtures were then treated with 300 μg of proteinase K at 65°C overnight. DNA was purified by the phenol-chloroform method. Purified DNA was dissolved in 150 μl of 10 mM Tris (pH 7.5). DNA was amplified by PCR using the following primers.

(3C assay Primer A-Forward : GGCGCCAGGCCGGGTCGA)

(3C assay Primer B-Reverse : TCGCGTCCCCGCTCCCCTATT)

(3C assay Primer C-Forward : CAGCCTCCTGATTGGCGGATAG)

(3C assay Primer D-Reverse : CCACCATCTTCCCACCCTCAG)

### Yeast two-hybrid screening

Yeast two-hybrid screening was carried out using the Matchmaker Gold Yeast Two-Hybrid system (Clontech/Takara, Mountain View, CA, USA). The C-terminal region of BRD2 (aa 450–802) was used as bait. Prey proteins were expressed from the universal human cDNA library (Cat. No. 630481;Clontech/Takara). Bait- and prey-transformed yeast (strain Y187) were mated, and the resultant diploids were cultured in DDO (SD-Leu/-Trp) medium to select for the presence of both plasmids. Subsequently, the diploids were cultured in DDO/X/A (SD-Leu/-Trp/X-α-gal/ABA) and QDO/X/A (SD-Leu/-Trp/-His/-Ade/X-α-gal/ABA) medium to select diploids exhibiting protein–protein interactions. Ten-fold serial dilutions were performed prior to colony plating in order to ensure that growth on the selective medium was dependent solely on *HIS3, ADE2, ABA*, and *MEL1* reporter gene expression. Selected colonies were subjected to DNA sequencing. Leu, leucine; Trp, tryptophan; His, histidine; Ade, adenine; SD, synthetic defined medium; ABA, aureobasidin A (antibiotic).

### Apoptosis assay by Annexin V staining

Cells were harvested and processed using the FITC–Annexin V Apoptosis Detection Kit I (BD Biosciences, San Jose, CA, USA) and analyzed by flow cytometry on a BD FACSCalibur machine (BD Biosciences).

### RNA-seq analysis

Isolated total RNA was processed for preparation of an RNA-seq library using the Illumina TruSeq Stranded mRNA Sample Preparation kit (Illumina, San Diego, CA, USA). Quality and size of libraries were assessed using the Agilent 2100 Bioanalyzer DNA kit (Agilent, Santa Clara, CA, USA). All libraries were quantified by qPCR using a CFX96 Real Time System (Bio-Rad, Hercules, CA, USA) and sequenced on NextSeq500 sequencers (Illumina). Sequencing adapters and low-quality bases in the raw reads were trimmed using the Cutadapt software. The cleaned high-quality reads were mapped to the human reference genome hg19 (https://genome.ucsc.edu) using STAR software. Genes differentially expressed between two selected biological conditions were identified by Cuffdiff in the Cufflinks package (http://cole-trapnell-lab.github.io/cufflinks/papers/).

### Quantification and Statistical analysis

For cell line studies, heatmaps were analyzed by the log-rank test using the PermutMatrix software ^36^. Gene clustering was analyzed using DAVID Bioinformatics Resources 6.8 ^34^.

